# Area requirements to safeguard Earth’s marine species

**DOI:** 10.1101/808790

**Authors:** Kendall R. Jones, Carissa Klein, Hedley S. Grantham, Hugh P. Possingham, Benjamin S. Halpern, Neil D. Burgess, Stuart H.M. Butchart, John G. Robinson, Naomi Kingston, James E.M. Watson

## Abstract

Despite global policy commitments to preserve Earth’s marine biodiversity, many species are in a state of decline. Using data on 22,885 marine species, we identify 8.5 million km^2^ of priority areas that complement existing areas of conservation and biodiversity importance. New conservation priorities are found in over half (56%) of all coastal nations, with key priority regions in the northwest Pacific Ocean and Atlantic Ocean. We identify where different conservation actions, ranging from marine protected areas to broader policy approaches, may best overcome anthropogenic threats to these areas. This analysis shows 26-41% of the ocean (depending on targets used for species representation) needs to be effectively conserved through a combination of site-based actions and wider policy responses to achieve global conservation and sustainable development agendas.

**One Sentence Summary:** At least 26% of the ocean needs a combination of site-based conservation and wider policy responses to achieve global conservation goals.

## Introduction

Biodiversity loss is one of the biggest environmental issues of our time (*1*). Human activities associated with agriculture, urbanisation, and natural resource extraction have led to large-scale habitat destruction and degradation, causing not only species declines and extinctions (*2*, *3*), but also the rapid erosion of intact ecosystems on land and in the sea (*4*, *5*). The disparity between increasing conservation efforts, including a doubling of the protected area estate in just two decades (*6*), and persistent biodiversity decline has led to a number of calls for more ambitious, science-based plans to halt biodiversity loss (*7*–*10*).

There is a clear scientific basis for substantially increasing area-based conservation efforts (*11*–*13*), and discussions around the generation of a new international framework for biodiversity conservation are now well underway (*14*). This framework will require not only the growth of strict, well-funded and well-managed protected areas, but also actions to manage the entire land/seascape to ensure conservation of biological diversity, sustainable use of its components, and fair and equitable sharing of its benefits (*7*, *15*). As a new, post 2020 biodiversity framework is currently being debated, it is crucial that baseline analyses provide the necessary detail on where and how conservation action is needed to safeguard biodiversity now. This is especially true in the ocean, where protected area coverage is substantially lower than on land (*6*), and existing conservation efforts are totally insufficient for the majority of marine species (*16*).

Here we identify global priorities for the expansion of conservation efforts, encompassing both site-based conservation action (including formal protected areas and other approaches) and broad-scale policy responses (e.g. addressing fisheries management) to secure marine species. We first evaluate how well ~23,000 marine species (fishes, mammals and invertebrates) are represented within current marine protected areas (MPAs), key biodiversity areas (KBAs; *17*), and the ocean’s remaining wilderness areas (*4*). We chose MPAs, KBAs and marine wilderness areas as the conservation baseline because all three currently play an essential role in safeguarding marine life, although only MPAs, as a governance structure, are actively managed. Well placed and resourced MPAs are critically important in stabilising or increasing species populations (*18*), maintaining coral cover (*19*), and generally have higher biomass than unprotected areas (*20*). Similarly, marine KBAs are sites of significance for the global persistence of biodiversity, supporting threatened or geographically restricted species/ecosystems, intact ecological communities, important biological processes (e.g. breeding aggregations), and/or having high irreplaceability (*17*). Marine wilderness areas are, by definition, the least anthropogenically impacted areas of the ocean, and so are mostly free of threats to biodiversity, at least for now (*4*). Wilderness areas, while not necessarily highly biodiverse, also often contain high genetic diversity, unique functional traits and endemic species, and much higher biomass than more anthropogenically impacted areas, so safeguarding them is critical in a time of human-forced climate change (*4*, *21*). Some KBAs and marine wilderness areas are contained within MPAs, or are seen as core priorities for future MPA expansion (*4*, *17*), and they are also important in informing areas that may be defined as “other effective area-based conservation measures” (OECMs: *22*) or managed through broad-scale policy approaches (e.g. fisheries restrictions). We hereafter refer to these areas (MPAs, KBAs and marine wilderness) as “areas of conservation or biodiversity importance”, as all offer accepted conservation benefits through direct protection (i.e. MPAs) or as areas of documented biodiversity importance (e.g. KBAs, marine wilderness).

Our analysis identifies species that have none of their range contained within areas of existing conservation or biodiversity importance, as well as those that do not meet various representation targets (we focus on the minimum target of 10% of total range covered, although other targets were explored). We then use an algorithm for solving integer linear programming problems (*23*) to identify additional conservation priorities to achieve coverage targets for each species while minimising the total area required. To assess the actions needed to conserve species within these areas, we then map the intensity of 15 damaging human activities across them, using the most comprehensive database of human stressors to the ocean (*24*). We distinguish between ocean-based stressors (e.g., fishing)-which can be managed with MPAs, OECMs, or other regulations concerning ocean activities (e.g. fishing, shipping) - and land-based stressors (e.g., nutrient runoff) which require terrestrial management. We ignore stressors where local actions have limited benefit (e.g. climate change). By doing this, we present an ecologically relevant, action-oriented plan to inform future marine conservation frameworks.

## Results

### Current species coverage

Using data on the global distribution of MPAs, we find that two-thirds of species (n = 15,149) meet a target of >10% range coverage by protected areas (Figure 1A;, *25*). Coverage levels vary considerably across marine taxa. Reptiles (n = 32) are the most well-covered group, with >90% of species meeting the 10% threshold, and all species having >2% of their range within MPAs (Figure 1A). In contrast, three percent of 3556 arthropod species have none of their range covered by protected areas, and only one-third of 117 mammal species reach the 10% threshold (Figure 1A). In total, 7736 species (33%) have <10% of their range covered by current MPAs. Around half of these species have under five percent of their range covered, and 216 species (~1%) have no part of their range within MPAs (Figure 1A).

**Fig. 1.**
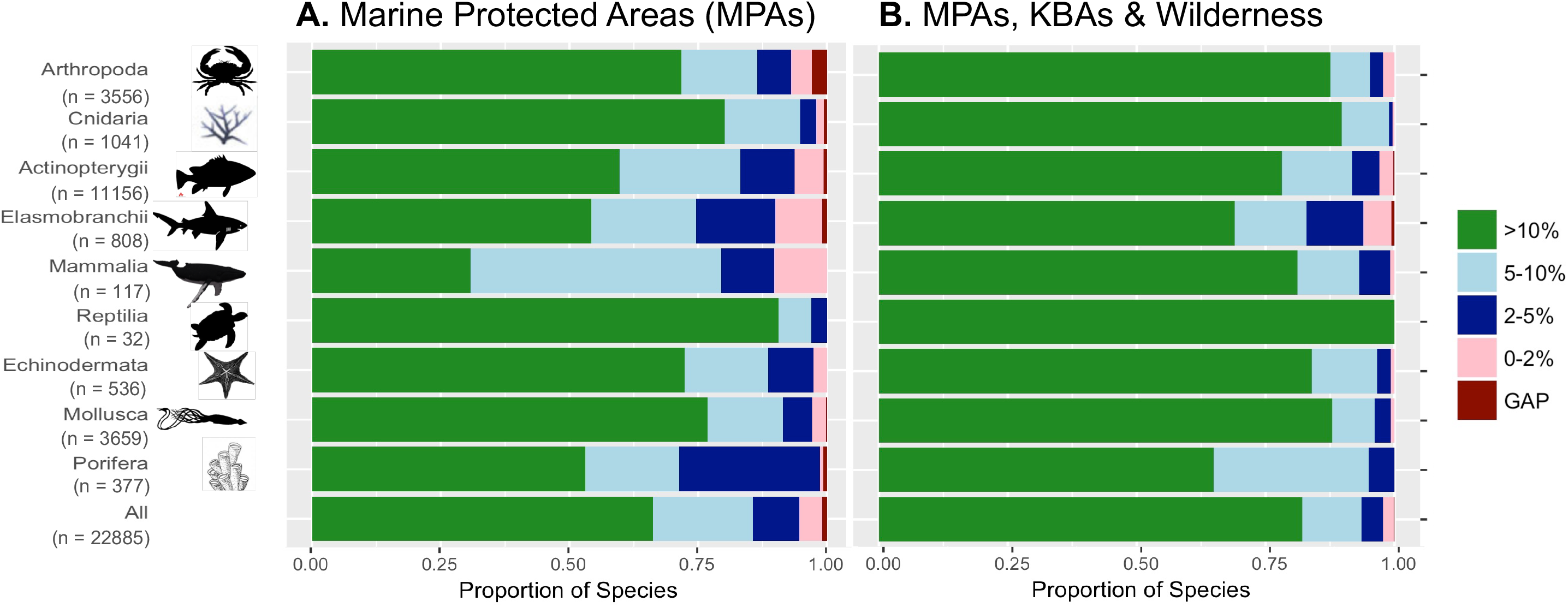
Protection levels across marine taxa. Percentage of marine species with 0% (dark red), 0–2% (pink), 2–5% (dark blue), 5–10% (light blue), and >10% (green) of their range overlapping with **A.** marine protected areas (MPAs), **B.** MPAs, key biodiversity areas (KBAs), and marine wilderness areas. Data are shown for all species (bottom) and species in the 6 largest phyla where the largest phyla (Chordata) is split into its 4 largest classes (Actinopterygii, Chondrichthyes, Mammalia, Reptilia).

We repeated this analysis to include all areas of conservation or biodiversity importance (MPAs, KBAs and marine wilderness areas), finding that species representation improved, with 82% of all species (n = 18,804) having >10% of their range inside MPAs, KBAs or marine wilderness (Figure 1B). Despite only 33 species (<0.1%) having no part of their range covered, there are still 4081 (18%) species with <10% coverage, and 500 (2%) species with <2% coverage (Figure 1B). Low coverage species (<2% coverage) are mostly found in the Atlantic Ocean, especially between Africa and South America, and also in the Pacific near China and Japan (Figure S1). Elasmobranchs (sharks and rays), and Porifera (sponges), are the least well covered phyla overall, with one-third of species having <10% coverage (Figure 1B). Under range-size based targets, 25.8% of species are adequately represented when considering only MPAs, and 66.1% of species are adequately represented when considering all areas of conservation or biodiversity importance.

### Global conservation priorities

We mapped global marine conservation priorities using integer linear programming (*23*) to identify where additional conservation responses, beyond effective conservation of MPAs, KBAs and marine wilderness areas, are needed to meet the representation targets for all species in the minimum total area. Because neither Aichi target 11 of the UN Convention on Biological Diversity (CBD) nor the UN Sustainable Development Goals (SDG) set specific targets for protecting individual marine species ranges, we set a minimum target of 10% of each species’ range (as both mechanisms aim to conserve at least 10% of the ocean, especially “areas of particular importance for biodiversity”; (*25*, *26*). However, we also explored a scenario which sets high targets (100%) for small-ranged species, lower targets (10%) for large-ranged species, and linearly scales targets between these values for medium-ranged species (see supplementary materials). We did not include Ecologically or Biologically Significant Marine Areas (EBSAs) because many are mapped at such large spatial scales that they are not useful as targets for area-based conservation without further refinement. We used a uniform proxy for cost data (area of conservation zones), rather than socio-economic data (e.g. fishing effort), to avoid inaccuracies and errors in socio-economic data biasing selection of planning units (*27*).

Representing 10% of all mapped species ranges would require 8.5 million km^2^ (~2.5% of the ocean) of new conservation priority areas in total, with just over half (55.4%, 4.7 million km^2^) located inside exclusive economic zones (EEZs; Figure 2; Figure S2). Combined with existing MPAs, KBAs, and marine wilderness, these areas cover 94.3 million km^2^ (26%) of the ocean (Figure 2). In comparison, expanding priority areas to meet species range-size based targets, which vary from 10% for large-ranged species to 100% for small-ranged species (*28*), would require 66 million km^2^ of new conservation priorities. When combined with existing areas of conservation and biodiversity importance, these expanded priorities would cover 152 million km^2^ (41%) of the ocean (Figure S3).

**Fig. 2.**
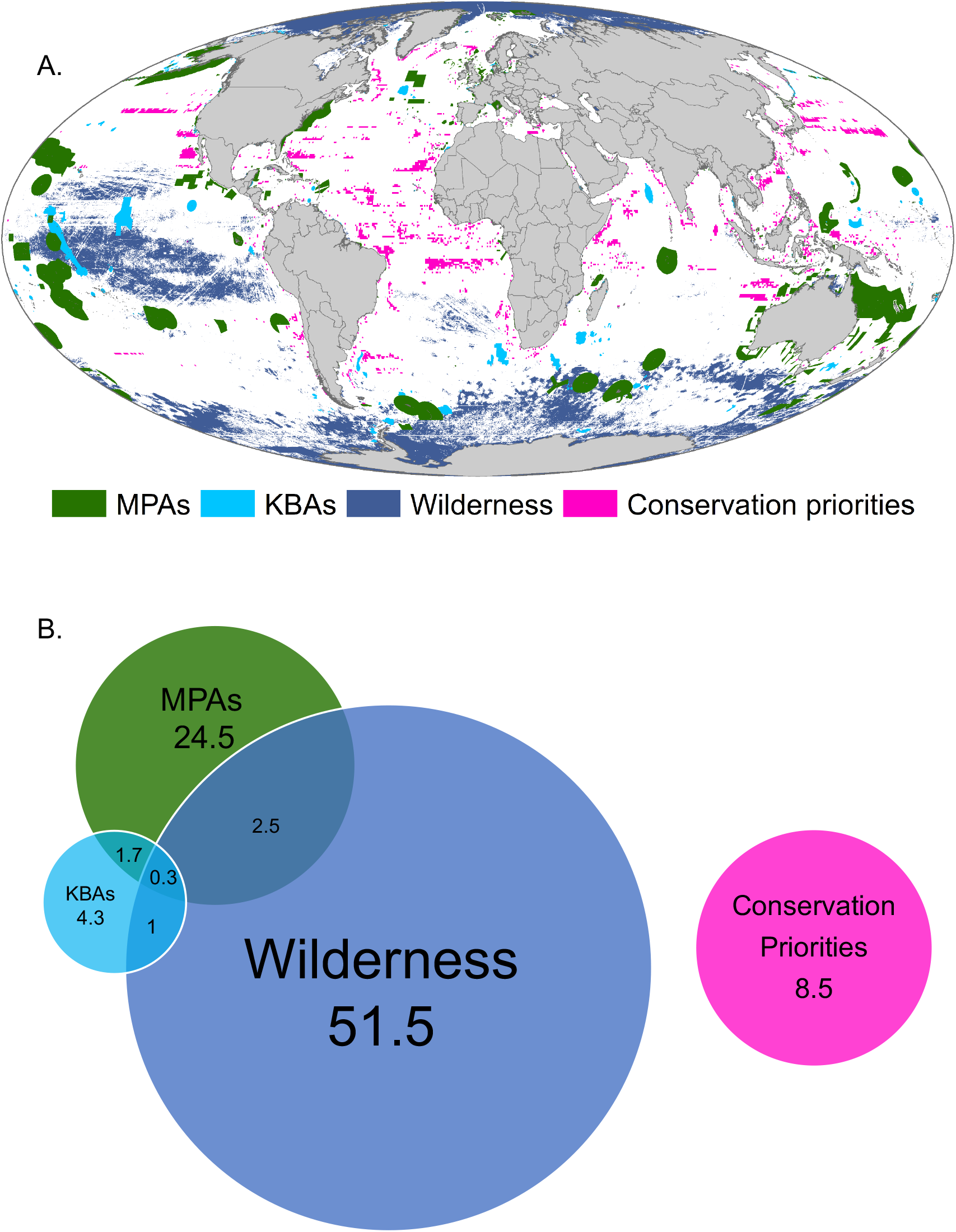
Location and size of conservation priorities. **A.** Minimum area required for conservation action to reach 10% coverage for approximately 23,000 marine species with known distributions, while accounting for existing marine protected areas (MPAs), key biodiversity areas (KBAs), and marine wilderness areas. **B.** Area of each mapped zone and their overlaps, in millions of km^2^.

New conservation priorities based on the 10% range target scenario are primarily located in places where there are few existing areas of conservation or biodiversity importance, and high concentrations of poorly represented species. Key regions for these priority areas include the Northern Pacific Ocean near China and Japan and the Atlantic Ocean between West Africa and the Americas (Figure S1). While we excluded EBSAs from our analysis, our new and existing priority areas (under the 10% range target scenario) overlap with 89% of individual EBSAs and cover 22% of total EBSA area.

Just over half (56%) of all coastal nations contain new priority areas (under the 10% range target scenario), although the amount within each country varies considerably (Figure 3). Of the new conservation priorities within EEZs, almost half are found in Asian and North American countries (Figure 3). Brazil has the largest area of new conservation priorities (452,000 km^2^), 64,000 km^2^ more than that of the next highest nation Indonesia (388,000 km^2^; Figure 3). Some nations with very large MPA estates still contain a substantial extent of new priority areas. For example, the United States has the largest MPA estate in the world (*6*), but the nation contains 168,000 km^2^ of new conservation priority areas (Figure 3), in part because it has the largest EEZ in the world, spanning three oceans. Timor-leste, The Bahamas, and the Chinese province of Taiwan have the largest area of conservation priorities relative to EEZ size, all having > 20% of their EEZ covered by new priority areas (Figure S2). Under a scenario using range-size based species representation targets, priority areas are found in almost all in coastal regions, throughout the Atlantic Ocean, and in the west Pacific Ocean (Figure S3)

**Fig 3.**
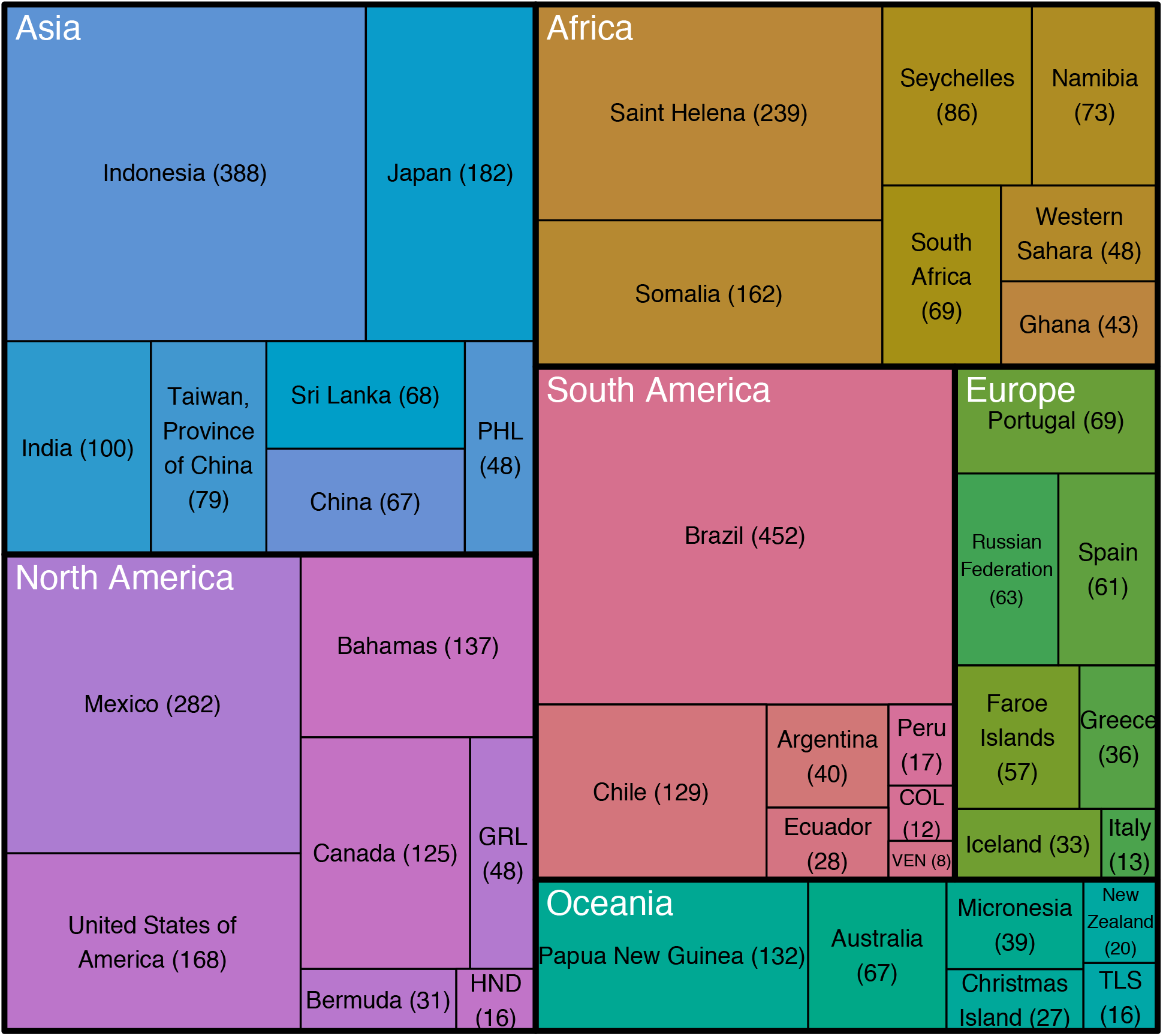
Size of conservation priority areas per continent. Area (thousands of km^2^) of new conservation priorities (under the 10% range target scenario) within Exclusive Economic Zones of the seven countries within each continent that contain the largest area of new conservation priorities. The size of each section is proportional to the area of conservation priorities within each continent and country. Antarctica is excluded as it is the territory of multiple nations. COL = Colombia, GRL = Greenland, HND = Honduras, PHL = Phillipines, TLS = Timor-leste, VEN = Venezuela.

### Assessing threats to priority areas

To assess threats to species across new conservation priority areas, we used the most comprehensive, globally consistent database on 19 human stressors to the marine environment (*24*). We excluded four climate stressors as they can only be effectively addressed through global action to reduce emissions, while here we focus on conservation actions that can be taken at the local to national scale. We classified the 15 remaining stressors based on whether they are ocean-based (e.g. fishing, commercial shipping) and can thus be managed with strict MPAs or other spatial regulations, or are land-based (e.g. nutrient runoff) and will require terrestrial actions such as land-use management to reduce runoff (Table S1).

Most new conservation priority areas (under the 10% range target scenario) are impacted primarily by ocean-based threats because the footprint of land-based pressures is constrained to near-coastal areas. Key areas of ocean-based threats to new conservation priorities include the East-China Sea and the North Sea off the Norwegian coast, which are both areas of intense industrial fishing activity (*29*; Figure 4 – blue colours). Ocean-based threats are generally lower in high-seas areas than near-shore priority areas, especially in the South Atlantic Ocean (Figure 4). While some coastal priority areas show very low levels of ocean-based threats, this may be partly due to a lack of data on fishing activity. For example, in Somalia it is estimated that illegal, unregulated and unreported fishing catch is almost three times higher than official estimates (*30*).

**Fig 4.**
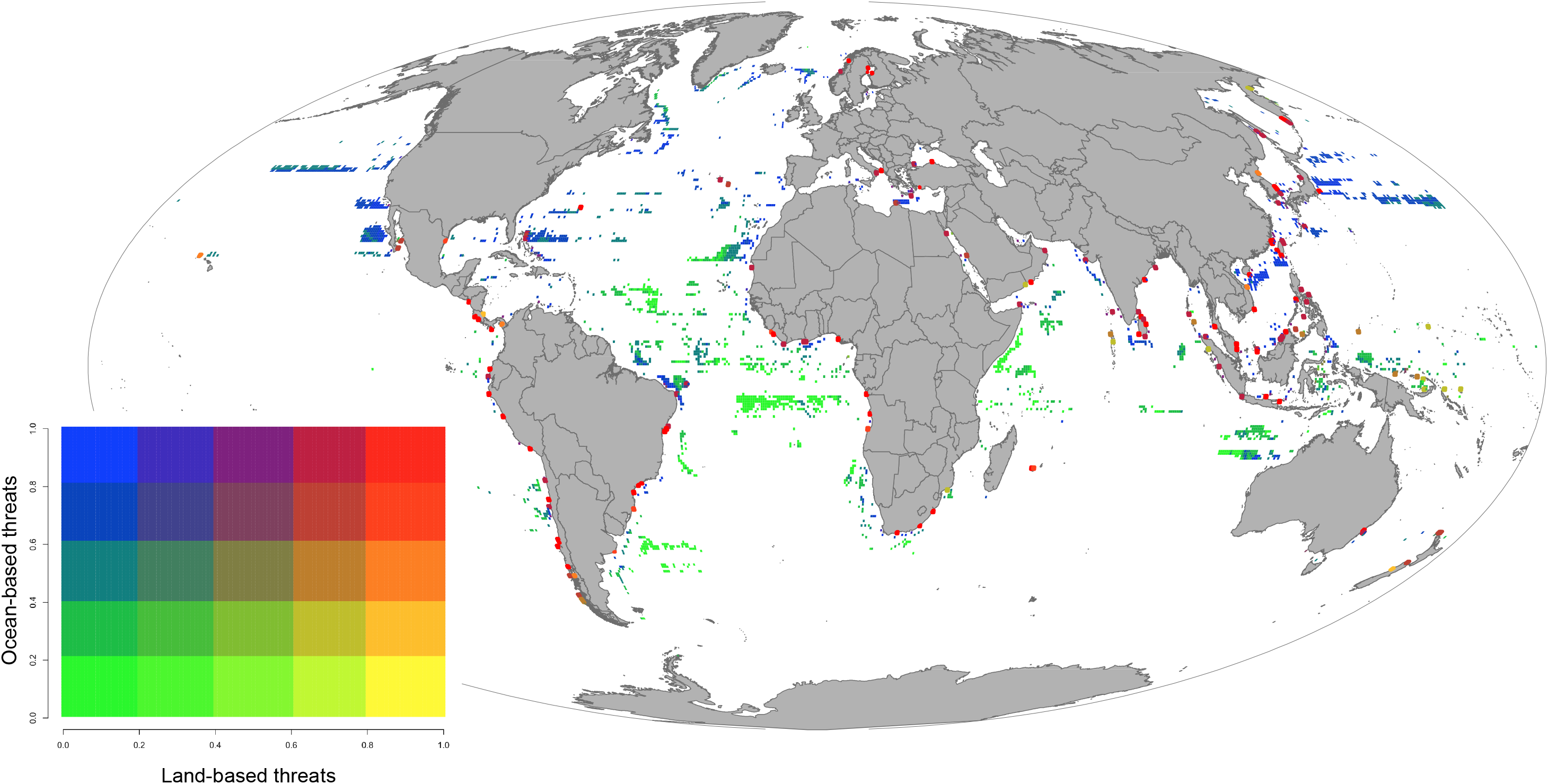
Threats to new priority areas for conservation. Spatial relationship between ocean-based threats (e.g. fishing, shipping; blue areas), and land-based threats (e.g. pollution runoff, nutrient runoff; yellow areas) across global priority areas for conservation (under the 10% range target scenario). Areas with high levels of ocean and land-based threats are shown in red, and those with low levels of ocean and land-based threat are shown in green. Boundaries of areas within the top two quintiles of land-based threat level (orange/red colors) have been enlarged to increase visibility.

A small number of new conservation priority areas (under the 10% range target scenario) are impacted by high levels of both ocean and land-based threats (Figure 4 – red colours). These impacts are highest in areas where high fishing activity coincides with high levels of agriculture and livestock grazing in very large nearby drainage basins, such as the Gulf of Mexico and the South China Sea. Many of these areas, such as the Indus river in Pakistan, have been previously identified as threat hotspots where coordinated management of land and ocean based impacts is vital (*31*). There are few new priority areas that are affected primarily by high levels of land-based threats, but exceptions include Southern Chile and parts of Indonesia (Figure 4 – yellow colours).

A substantial proportion of new conservation priorities (under the 10% range target scenario) are currently facing relatively low overall threat (Figure 4 – green colours). It is crucial to prevent threats from expanding into these areas, as low-pressure habitat holds levels of biodiversity unparalleled in areas of higher impact (*32*). Monitoring is essential in these areas, but as they are often remote and located beyond national jurisdiction this can be difficult. However, with advances in remote vessel monitoring technology, such as Global Fishing Watch (*29*), this is becoming increasingly possible.

## Discussion

Future global strategies to address biodiversity loss will require rapid action to secure imperiled species and ecosystems, combined with long-term approaches to sustainably manage the ocean in its entirety (*8*). We show that effective conservation of an additional 8.5 million km^2^ - alongside improved management of existing MPAs, and proactive conservation of KBAs and marine wilderness (which cover ~60 million km^2^) - could achieve a minimum representation target of 10% for all mapped marine species. If species range-size based targets are used, an additional 15% of the ocean (~57 million km^2^) would require conservation. Regardless of the target used, our findings echo previous analyses showing that meeting global conservation targets for all mapped terrestrial species ranges will require large increases in the total area under conservation (*33*).

Our findings, showing that at least 26% of the ocean needs to be conserved are similar to recent calls from CBD parties and observers for a post-2020 conservation framework that ensures at least 30% of Earth is covered by effectively managed PAs and OECMs (*34*, *35*).

However, we are unable to quantify the proportion of priority areas that require conserving through site-based approaches (i.e. PAs and OECMs) versus broad-scale policy responses. Furthernore, our figure of 26% is likely a significant underestimate of the total area required. Beyond the fact that this result was based on a minimalistic 10% species representation target, our species data notably exludes all marine birds and represents a tiny fraction all marine species (*36*). We also do not consider the fact that species are changing their distributions in response to climate change (*37*), and assume that conservation of all areas within a species’ range contribute equally towards its conservation, ignoring areas important for different life-history stages (e.g., breeding grounds, feeding areas, migration routes). Furthermore, recently adopted KBA criteria (*17*) have yet to be applied to many taxa and ecosystems, so the marine KBA network will increase in the near future.

The benefits of marine conservation actions (e.g. MPAs, OECMs) are clearly commensurate with good design, adequate resourcing, fair governance and equitable management, which are lacking in many countries (*38*). However, our results assume that all MPAs are effective in stopping threats to biodiversity, likely vastly overestimating their conservation impact (*39*). For effective conservation, MPAs should conform to the IUCN’s global standards for conservation success (*34*), and as such, it is timely to consider revising the status of some areas currently considered MPAs (e.g. removing areas which are managed for fishing and have little or no conservation benefit). Furthermore, it is crucial to recognise that well-managed MPAs are only part of a suite of management options necessary to maintain ocean health (*34*), and that they must be combined with OECMS, land-based actions, and broad-scale approaches leading to improved management of the ocean beyond the protected area estate (e.g. fishery sustainability reporting, mandatory environmental impact asessments for extractive projects). These broad-scale policy approaches will often be better suited for wilderness conservation than MPAs, given the size and remoteness of many wilderness areas.

It is important to target management actions to the threats facing species in conservation priorities. Areas affected primarily by ocean-based threats are priorities for MPA designation, OECMs, or broad-scale policy responses such as strictly enforced fisheries regulations. However, many of these areas also support highly productive fisheries, meaning regulations can fail in the face of intense opposition from fishers (*40*). Overcoming these challenges will require identifying which species and ecosystems are most vulnerable to ocean-based impacts, and thus require strict protection to prevent extinctions, and also identifying where conservation outcomes can be achieved while allowing sustainable resource extraction. It is also crucial to recognise that the priority areas we identify require effective conservation, which does not necessarily equate to strict protection in MPAs. In many cases, management in OECMs or through broad-scale policy mechanisms may be more appropriate, particularly for extensive marine wilderness areas where site-based conservation may be inappropriate. In areas where land-based stressors play a dominant role in determining ecosystem condition, MPAs will have little benefit unless the adjacent land is also managed for conservation (*31*). This may involve triaging areas where high levels of land-based stressors make achieving conservation outcomes unlikely or prohibitively expensive. In other areas, species-targeted gear restrictions might be preferable to MPAs, especially for pelagic megafauna with wide distributions, or for species that are only threatened by a single fishery (*41*).

While addressing land and ocean-based threats is important in the immediate term, conservation strategies must also look forward to assess the future risk posed by human-forced climate change (*42*). Local conservation actions are unable to stop or reverse the impacts of climate change, but there are many actions that can increase the ability of biodiversity to adapt to a changing climate. For example, there is mixed evidence for MPAs enhancing recovery and resilience of degraded coral reefs (*43*), and reducing land-based stressors may increase reef resilience to climate change (*44*). However, most studies on how conservation efforts influence climate change resilience focus on coral reef ecosystems, so further research on other coastal ecosystems (e.g. seagrass, kelp forests) and pelagic systems is vital. It is also important to recognize and plan for the impacts of human responses to climate change, which include shifting fishing effort to track fish stocks, or building seawalls to prevent sea-level ris (*42*).

Because over 46% of priority areas are located in the high seas, developing and implementing conservation actions in these areas will be crucial for future conservation agreements. Conservation action in the high seas is legally challenging and has so far been limited, with only 1.18% currently protected (*6*). However, the need for high-seas management is also being increasingly recognised by the international community, with the UN currently negotiating a legally binding high-seas conservation treaty to be established under the existing Law of the Sea Convention (*45*). Legal options for conservation under such an agreement are still being debated, but it will likely provide an opportunity to increase the use of area based conservation management tools (e.g. MPAs, fishing restrictions), and to mandate environmental impact assessments for all activities occurring in the high-seas (*45*).

Given the difficulties in establishing MPAs in the high seas, another option is to use existing international and regional agreements to achieve conservation goals. For example, Regional Fisheries Management Organisations (RFMOs) — international organizations formed by countries to manage shared fishing interests in a certain area — are already used to set catch and fishing effort limits (*46*). In some areas, RFMOs have even been used to protect vulnerable marine ecosystems from bottom-trawl fishing (*46*), so use of these powers to create more high seas conservation areas is certainly feasible (*45*). Alternatively, given that 54% of high seas fishing would be unprofitable without government subsidies, subsidy reform could also act as a useful management tool for high seas fisheries (*47*).

With the 2020 deadline for achieving global conservation targets under the CBD fast approaching, this work highlights new priorities for conservation action to fulfill current targets. Our work shows that effective conservation of an additional 8.5 million km^2^ – alongside improved management of existing MPAs, and proactive conservation of KBAs and marine wilderness - could secure 10% of all mapped marine species ranges. This effort would lead to considerable progress towards the life below water goal (SDG14) of the UN Sustainable Development Goals (*26*). Our analysis presents a realistic, ecologically relevant baseline for the conservation community to strive towards, and if combined with an agenda focused on improved management of the ocean beyond the priorities identified here, represents the start of a bold plan for the future of marine conservation.

### Materials and Methods

All spatial data described below were processed using ESRI ArcGIS v10.5 in Mollweide equal-area projection. All prioritisation analyses were conducted using R statistical software 3.3.

### Gap analysis

Data on the global distribution of protected areas (PAs) were obtained from the 2017 World Database on Protected Areas (*48*). Following similar global PA studies (*49*) we extracted PAs from the WDPA database by selecting those areas that had a status of “designated”, “inscribed”, or “established”, and were not designated as UNESCO Man and Biosphere Reserves. We included only PAs with detailed geographic information in the database, excluding those represented as a point only. We then used a layer delineating global coastline to identify marine PAs (MPAs) by clipping PA polygons to only include those which have some overlap with marine area (http://datadryad.org/resource/doi:10.5061/dryad.6gb90.2). EBSA data was taken from https://www.cbd.int/ebsa/ebsas.

Data on Key Biodiversity Areas (KBAs) were obtained from the 2016 release of the World Database of Key Biodiversity Areas (http://www.keybiodiversityareas.org/). We used a layer of terrestrial country boundaries to clip KBA polygons to only include those which overlap with marine area http://datadryad.org/resource/doi:10.5061/dryad.6gb90.2). We used previously identified data on marine wilderness areas (*4*), which were mapped by identifying areas that have both little to no impact across 15 human stressors to the marine environment (excluding 4 climate stressors), and also a low combined impact from 19 human stressors including climate change stressors. To avoid double counting areas that are covered by MPAs, KBAs, and marine wilderness, we merged these three layers and dissolved areas where they overlapped.

2015 data on marine biodiversity was obtained from Aquamaps (*36*), a species distribution modelling tool that correlates known species occurrence points with environmental data (e.g. temperature, salinity) to produce standardised global range maps for 22,885 aquatic species. This is the most comprehensive and highest resolution data available on the distribution of marine biodiversity globally, and includes Animalia (fishes, marine mammals, and invertebrates), Plantae (fleshy algae, seagrass), Chromista (calcifying algae) and Protozoa. The species distribution maps predict relative probabilities of species occurrence (ranging from 0.00–1.00) at a resolution of 0.5-degree cells. It is assumed that the preferred range is where probability is 1, outside the range limits is where probability is 0, and between these two thresholds the relative environmental suitability decreases linearly. As there is no recommended threshold to use, we follow previous studies and report on results using probability threshold of 0.5 or greater (*16*). We did explore other probability thresholds and found that the results varied very little (see supplementary materials).

To assess coverage of marine species distributions in MPAs, KBAs and wilderness areas, we determined the proportion of protected area (MPA, KBA and wilderness) in each 0.5-degree cell. As we do not know the exact distribution of species within each cell, we assumed that the area of a species’ range represented in protected areas was equal to the protected area coverage for grid cells that species was present in.

### Spatial prioritisation analysis

We used integer linear programming to identify spatial priorities that meet a percentage coverage target for each of the 22,885 Aquamaps species, while accounting for the level of protection in existing MPAs, KBAs and wilderness, and minimizing the total cost of selected cells, with area as the cost, following previous studies (*23*). This is frequently referred to as the minimum-set problem in spatial conservation planning. We used the software package Gurobi (version 5.6.2) to find solutions to this minimum-set problem. Gurobi is proprietary software that uses several algorithms, including simplex and branch and bound algorithms, to solve linear programming problems and is guaranteed of finding optimal solutions given enough time. We set Gurobi to achieve a solution within 0.5% of the optimum (i.e. when the current solution was within 0.005 times the guaranteed lower boundary of the optimal solution). The optimal solution is that which achieves the coverage target with the lowest possible cost. We explored 3 different coverage targets: 10% of species range (reported in main text), 20% of species range, and a set of targets that varied depending on the total range size of each species (reported in supplemental materials).

We used 0.5-degree cells as our planning units (areas which can be selected or not selected for conservation), as this is the same scale as our species distribution data. We extracted all planning units containing species distribution records from Aquamaps (n = 178,234) and assigned each planning unit a cost value equal to the area of the cell that is not covered by an MPA, KBA or marine wilderness area. Thus, the cost value reflects the additional area per cell which requires management if selected for conservation.

### Assessing threats facing priority areas

We assessed the anthropogenic threats facing priority areas using normalized data on cumulative human impact to marine ecosystems (*24*). This threat database includes 19 individual human stressors, but we excluded four climate change stressors. We then categorized threats as ocean-based or land-based, depending on their origin (see Table S1 for full list). Ocean-based threats have clear marine origins, such as fishing and shipping, can therefore potentially be managed through effective MPAs of other ocean-use regulations, whereas land-based threats (e.g. nutrient runoff, coastal armouring) originate on land and will require land-management to address. All measures of fishing pressure, shipping (shipping lanes and ship-based pollution) and ocean structures (e.g. oil rigs) were considered as ‘ocean-based’ in our analysis, while all other threats were considered land-based. We summed the values for each individual stressor layer within the ocean-based and land-based stressor groups, to give final ocean-based and land-based human impact values. Using this information, we used the zonal statistics tool in ArcMap 10.5 to calculate the mean level of ocean and land-based threat within each planning unit selected as a priority area in our spatial prioritisation analysis.

### Sensitivity Analyses

The Aquamaps species distribution maps we used for our gap analysis and spatial prioritisation predict relative probabilities of species occurrence (ranging from 0.00–1.00) at a resolution of 0.5-degree cells. It is assumed that the preferred range is where probability is 1, outside the range limits is where probability is 0, and between these two thresholds the relative environmental suitability decreases linearly.To test the sensitivity of our results to the probability threshold used to determine species distributions within each 0.5-degree cell, we repeated our gap analysis using 4 probability thresholds: 0.25, 0.5, 0.75 and 1 (results presented in the main text use 0.5).

When using various probability thresholds to determine species distributions, the number of species within each coverage group (e.g. no coverage, 0-2% coverage etc.) varied by less than 1% across all probability thresholds tested (Table S1), and thus our results are not sensitive to species distribution modelling uncertainties. Furthermore, previous studies using Aquamaps data found that varying probability thresholds makes very little difference to global scale analyses (*16*).

To test the sensitivity of our results to the species representation target used, we explored 3 targets in our spatial prioritisation analyses, all of which aimed to minimse the total area of selected planning units: 10% of species range, 20% of species range, and range-size based targets that vary depending on the total range size of each species. For the latter case, we set a 100% coverage target for species with ranges < 10,000 km^2^, while for wide-ranging species (> 390,000 km^2^) the target was reduced to 10% coverage, and where geographic range size was intermediate between these extremes, the target was log-linearly interpolated. The 390,000 km^2^ threshold is arbitrary, but it follows previous studies (*28*) and corresponds to roughly one-third of all species analysed. For each set of targets, we set Gurobi to achieve a solution within 0.5% of the optimum (i.e. when the current solution was within 0.005 times the guaranteed lower boundary of the optimal solution). The optimal solution is that which achieves the coverage target with the lowest possible cost. As such, the prioritisations conducted in this sensitivity analysis identify sets of planning units that meet each species representation target (10% of all species ranges, 20% of all species ranges, and species range-size based targets) in the least possible area.

When using various species range targets in our spatial prioritization analysis, there was a high level of overlap between selected planning units, although the total area of priorities changed substantially. More than half of all the planning units selected when using a 10% coverage target were also selected when using a 20% target (Table S2). As expected, considerably more area was required to meet the range-size based targets, although 56% of planning units selected under the range-size based targets were also selected when using a 10% target. This suggests our priority areas are robust to different target setting approaches, as over 50% of planning units are always selected, regardless of the specific species representation target used. As such, future conservation agreements and priority setting exercises, which may use representation targets different to the 10% coverage target we report on in the main text, can efficiently build on the priority areas we identify.

### Bird range overlap

To explore how our conservation priority areas overlap with seabird ranges, we obtained data on the distribution of birds from Birdlife International (http://datazone.birdlife.org/species/requestdis). We extracted all birds classified as seabirds, and calculated the area of overlap between each seabird species range and our conservation priority areas. We found that our priority areas overlap with 67.4% (n= 247) of all seabird ranges, and cover 12.2% of individual species range area on average. In 42.8% (n= 157) of species, our priority areas cover >10% of their range.

## Acknowledgements

We would like to thank Stacy D. Jupiter, Emily S. Darling, Dan Lafolley and Enric Sala for providing constructive feedback on this study.

## Author contributions

K.R.J., C.J.K, H.P.P, and J.E.M.W. designed the study. K.R.J. carried out the analyses. All authors wrote and revised the manuscript.

## Data and materials availability

All data is available in the main text or the supplementary materials.

---

**Figure. S1.**
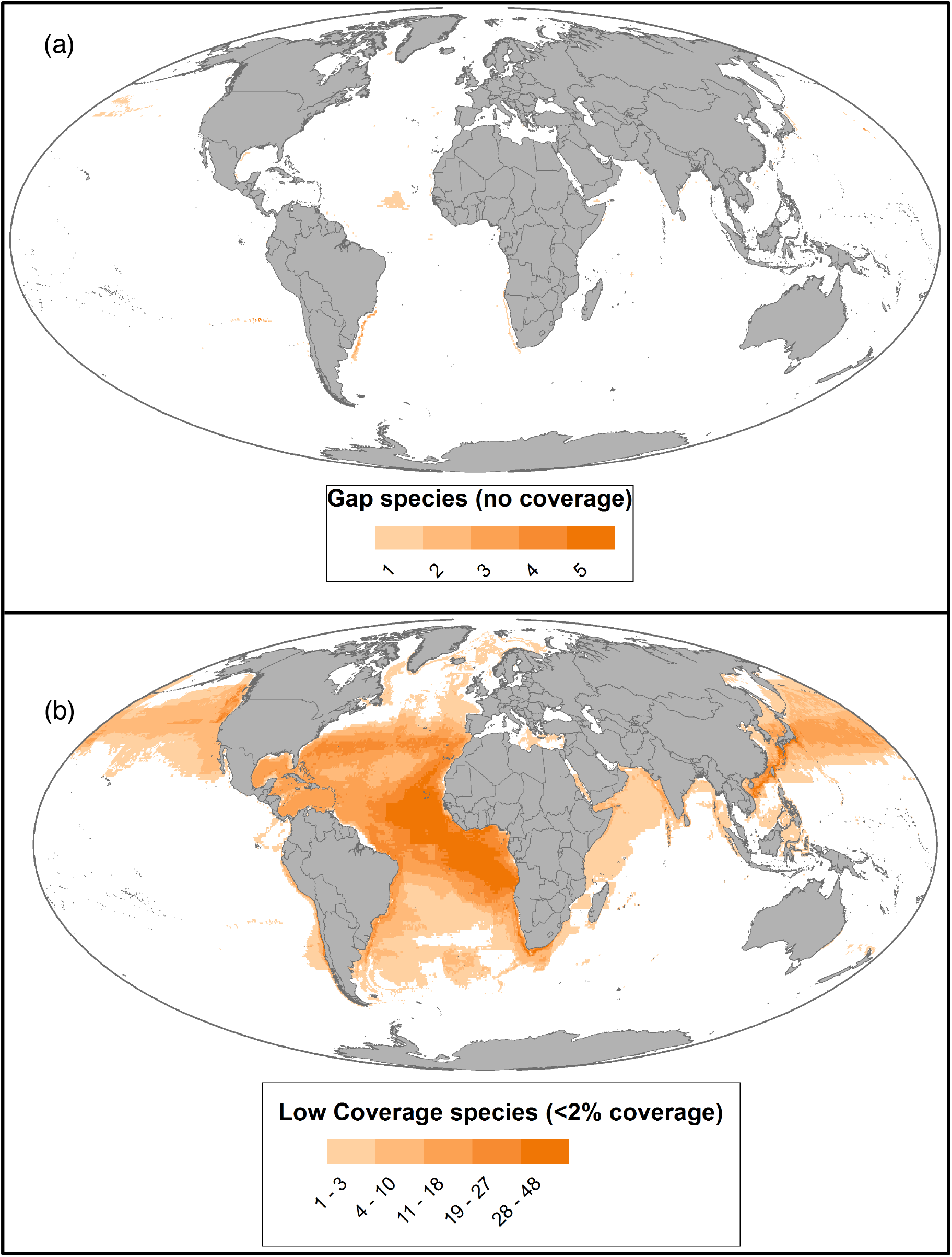
Density map of gap species (no range represented in MPAs, KBAs or marine wilderness) **(a)** and very low coverage species (<2% of range represented in MPAs, KBAs or marine wilderness) **(b)**.

**Figure. S2.**
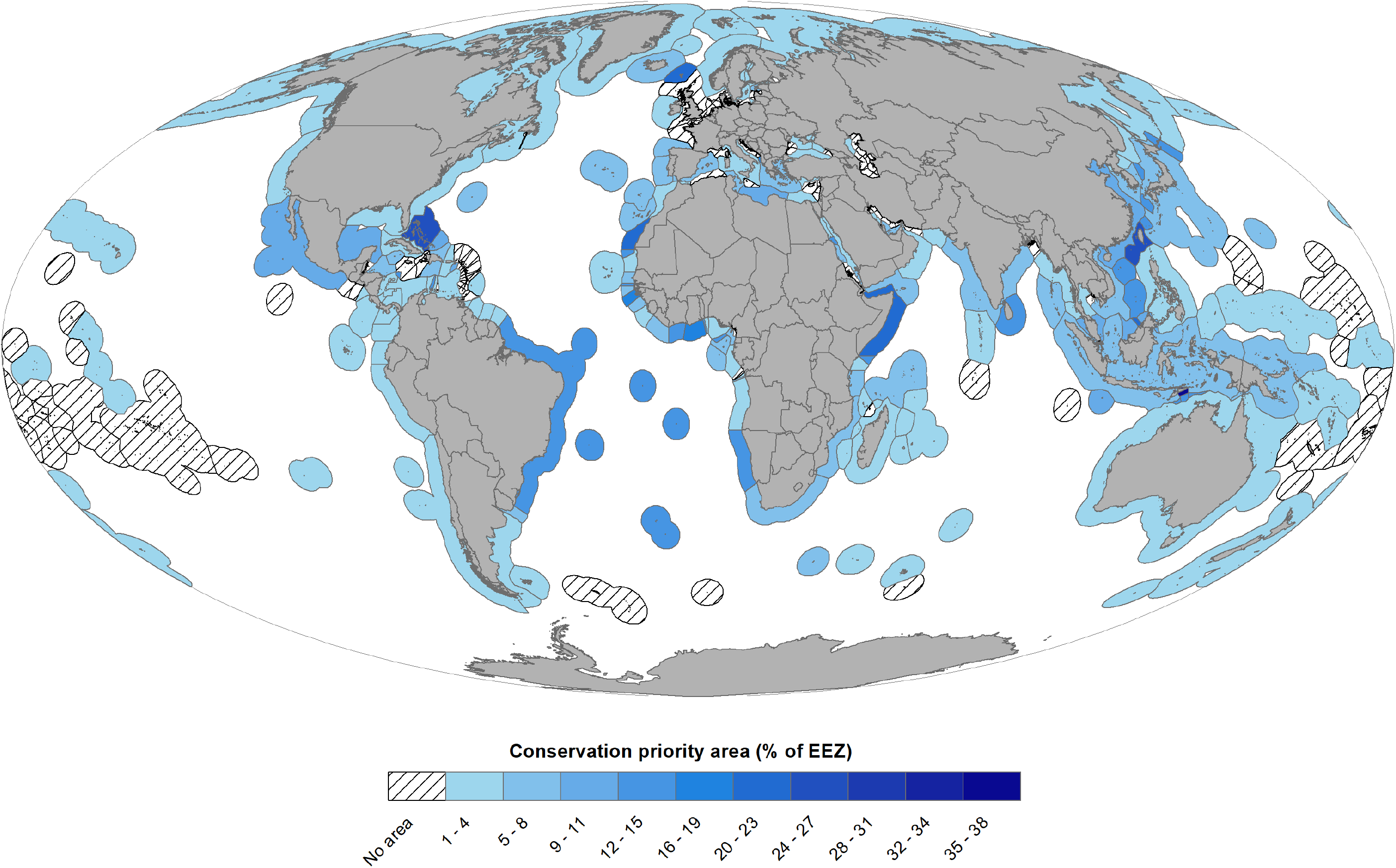
Area (% of EEZ) of new conservation priorities within Exclusive Economic Zones. Hatched areas contain no newly identified conservation priorities.

**Fig. S3.**
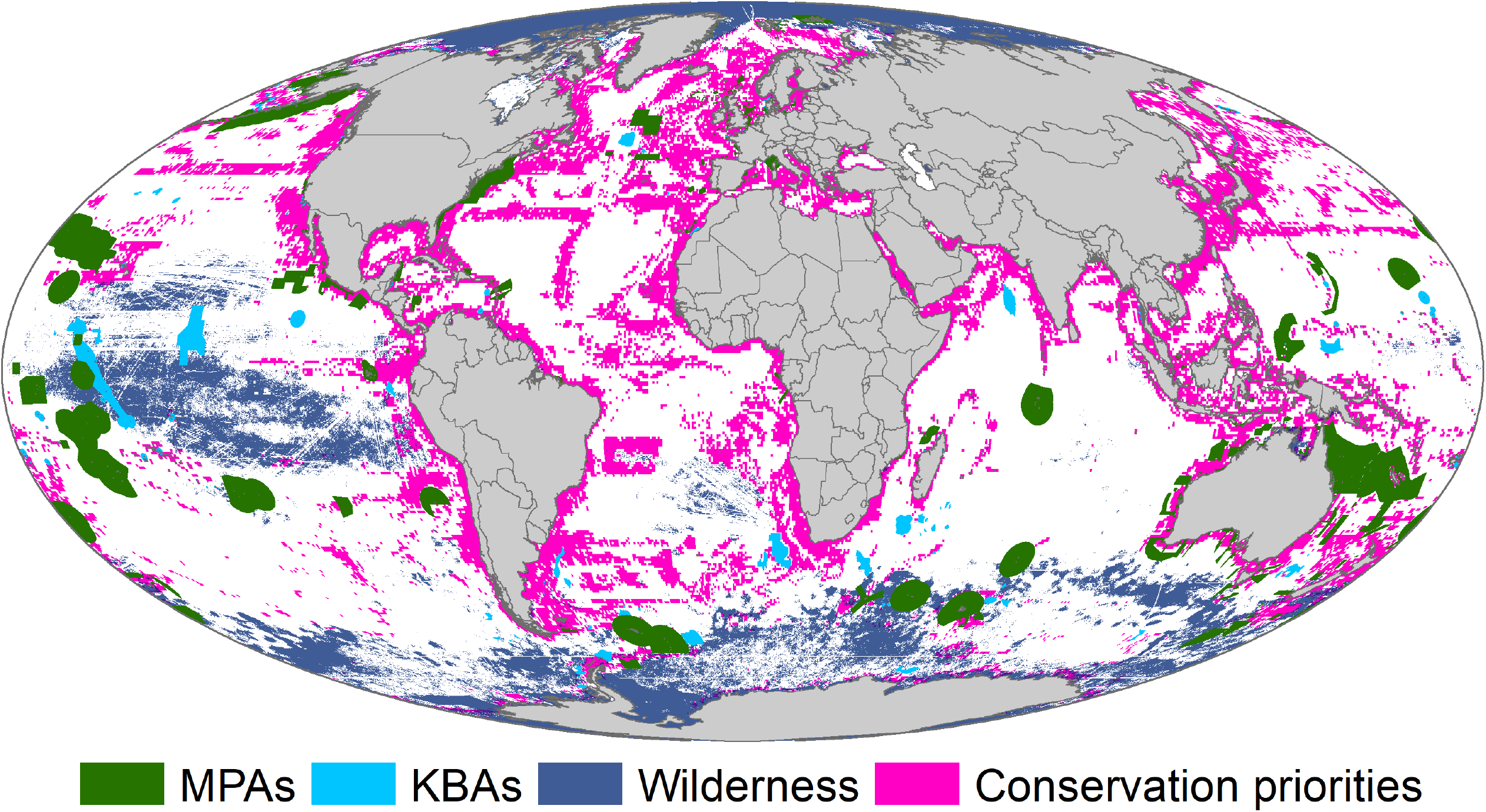
Minimum area required for conservation action to reach range-size based target coverage for approximately 23,000 marine species with known distributions, while accounting for existing marine protected areas (MPAs), key biodiversity areas (KBAs), and marine wilderness areas. Targets range from 10% for large-ranged species to 100% for small-ranged species.

**Table S1.**
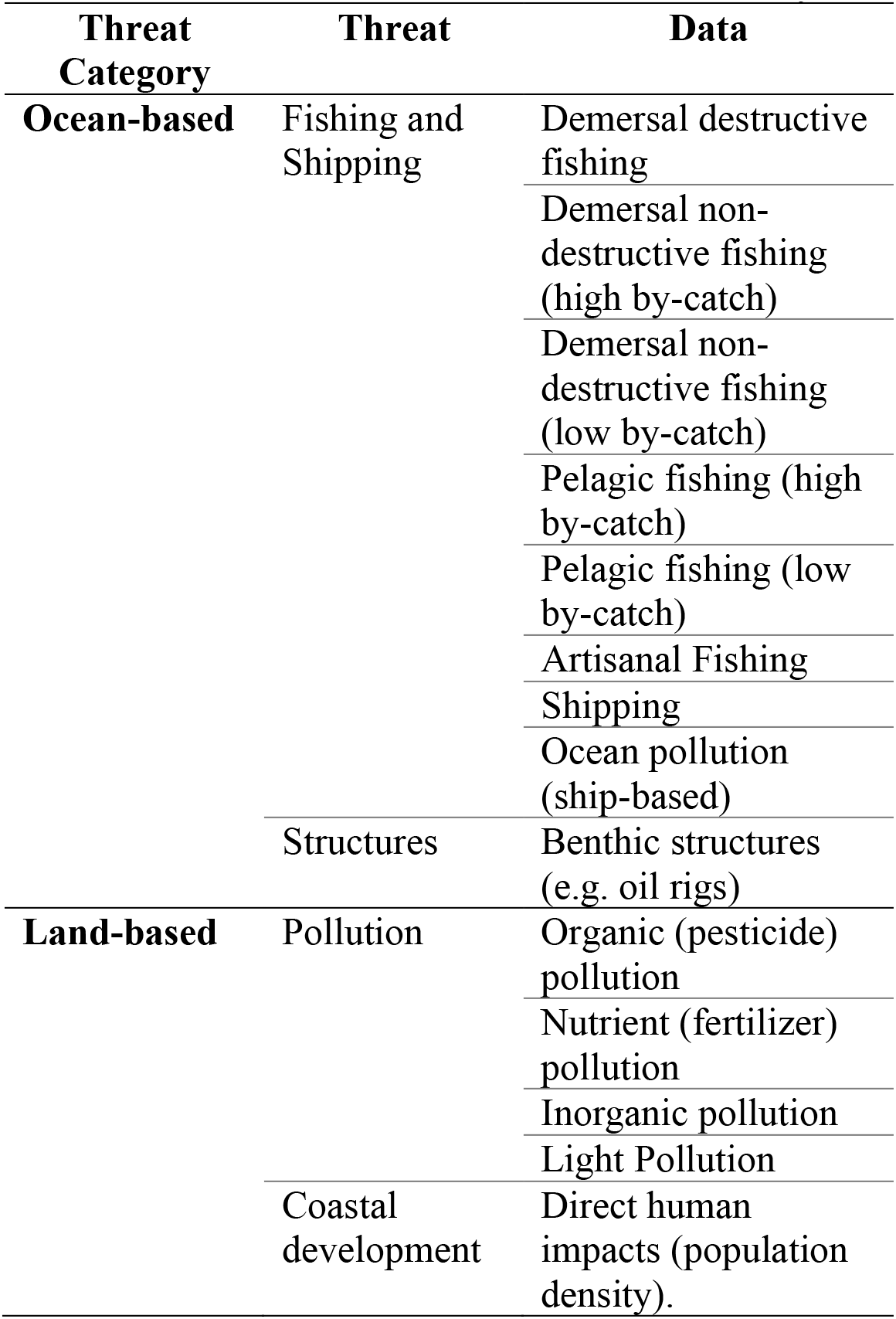
Classification of threats based on whether they are ocean-based or land-based (*24*).

**Table S2.** Proportion of marine species with 0% (gap), 0–2%, 2–5%, 5–10%, and >10% of their range overlapping with marine protected areas (IUCN I-VI), key biodiversity areas, and marine wilderness areas, for species probability thresholds ranging from 0.25–1.

| *Aquamaps probability threshold* | *Gap (no coverage)* | *Covered 0-2%* | *Covered 2-5%* | *Covered 5-10%* | *Covered >10%* |
| --- | --- | --- | --- | --- | --- |
| 0.25 | 0.1% | 1.8% | 4.0% | 11.3% | 82.8% |
| 0.5 | 0.1% | 2.0% | 4.2% | 11.4% | 82.2% |
| 0.75 | 0.2% | 2.5% | 4.5% | 11.7% | 81.1% |
| 1 | 0.1% | 1.8% | 4.0% | 11.3% | 82.8% |

**Table S3.**
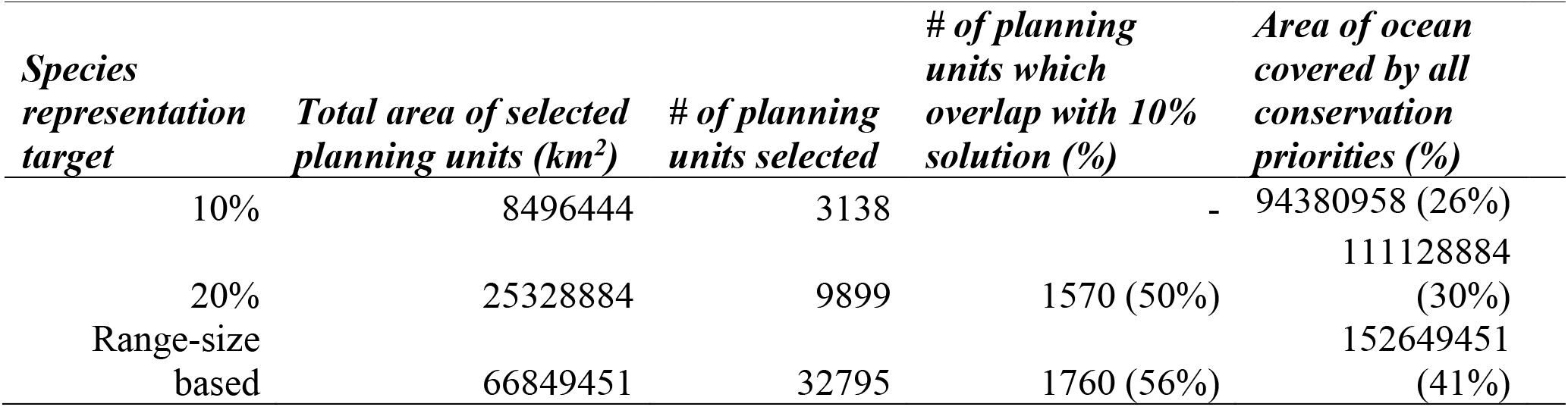
Total area, number of selected planning units, and the number of planning units which overlap with those selected when using a 10% coverage target, for two alternative coverage targets: 20% of all species ranges, and targets which vary depending on the total range size of species (see supplementary materials).

